# Stout camphor tree genome fills gaps in understanding of flowering plant genome and gene family evolution

**DOI:** 10.1101/371112

**Authors:** Shu-Miaw Chaw, Yu-Ching Liu, Han-Yu Wang, Yu-Wei Wu, Chan-Yi Ivy Lin, Chung-Shien Wu, Huei-Mien Ke, Lo-Yu Chang, Chih-Yao Hsu, Hui-Ting Yang, Edi Sudianto, Ming-Hung Hsu, Kun-Pin Wu, Ning-Ni Wang, Jim Leebens-Mack, Isheng. J. Tsai

## Abstract

We present reference-quality genome assembly and annotation for the stout camphor tree (SCT; *Cinnamomum kanehirae* [Laurales, Lauraceae]), the first sequenced member of the Magnoliidae comprising four orders (Laurales, Magnoliales, Canellales, and Piperales) and over 9,000 species. Phylogenomic analysis of 13 representative seed plant genomes indicates that magnoliid and eudicot lineages share more recent common ancestry relative to monocots. Two whole genome duplication events were inferred within the magnoliid lineage, one before divergence of Laurales and Magnoliales and the other within the Lauraceae. Small scale segmental duplications and tandem duplications also contributed to innovation in the evolutionary history of *Cinnamomum*. For example, expansion of terpenoid synthase subfamilies within the Laurales spawned the diversity of *Cinnamomum* monoterpenes and sesquiterpenes.

## Introduction

Aromatic medicinal plants have long been utilized as spices or curative agents throughout human history. In particular, many commercial essential oils are derived from flowering plants in the tree genus *Cinnamomum* L. (Lauraceae)^1-3^. For example, camphor, a bicyclic monoterpene ketone (C_10_H_16_O) that can be obtained from many members of this genus, has important industrial and pharmaceutical applications^4^. *Cinnamomum* includes approximately 250 species of evergreen aromatic trees belonging to Lauraceae (laurel family), which is an economically and ecologically important family that includes 2,850 species distributed mainly in tropical and subtropical regions of Asia and South America^5^. Among them, avocado (*Persea americana*), bay laurel (*Laurus nobilis*), camphor tree or camphor laurel (*C. camphora*), cassia (*C. cassia*), and cinnamon (including several *C.* spp.) are important spice and fruit species. Lauraceae has traditionally been classified as one of the seven families of Laurales, which together with Canellales, Piperales and Magnoliales constitute the Magnoliidae (“magnoliids” informally).

The magnoliids, containing about 9,000 species, are characterized by 3-merous flowers with diverse volatile secondary compounds, 1-pored pollen, and insect-pollination^6^. Many magnoliids – such as custard apple (*Annonaceae*), nutmeg (*Myristica*), black pepper (*Piper nigrum*), magnolia, and tulip tree (*Liriodendron tulipifera*) – produce economically important fruits, spices, essential oils, drugs, perfumes, timber, and horticultural ornamentals. The phylogenetic position of magnoliids, however, has been uncertain. Further, there are also unresolved questions about genome evolution within the Magnoliidae. Analysis of transcriptome sequences has implicated two rounds of genome duplication in the ancestry of *Persea* (Lauraceae) and one in the ancestry of *Liriodendron* (Magnoliaceae)^7^, but the relative timing of these events remains ambiguous.

*Cinnamomum kanehirae,* commonly known as the stout camphor tree (SCT), a name referring to its bulky, tall and strong trunk, is endemic to Taiwan and under threat of extinction. It has a restricted distribution in broadleaved forests in an elevational band between 450 and 1,200 meters^8^. *Cinnamomum,* including SCT and six congeneric species contributed to Taiwan’s position as the largest producer and exporter of camphor in the 19^th^ century, and its value was further enhanced due to its valuable wood, with trunks exhibiting the largest diameters among flowering plants of Taiwan, and aromatic, decay-resistance attributed to the essential oil D-terpinenol^9^. *Antrodia cinnamomea,* a parasitic fungus that infects the trunks of SCT causing heart rot^10^. The fungus produces several medicinal triterpenoids that impede the growth of liver cancer cells^10,11^ and act as antioxidants that protect against atherosclerosis^12^. Due to intensive deforestation in the past half century, followed by poor seed germination and illegal logging to cultivate the fungus, natural populations of SCT are fragmented and threatened^13,14^.

Here we report a chromosome-level genome assembly of SCT. Comparative analyses of the SCT genome with those of 10 other angiosperms and two gymnosperms (ginkgo and Norway spruce) allow us to resolve the phylogenetic position of the magnoliids and shed new light on flowering plant genome evolution. Several gene families appear to be uniquely expanded in the SCT lineage, including the terpenoid synthase superfamily. Terpenoids play vital primary roles as photosynthetic pigments (carotenoids), electron carriers (plastoquinone and ubiquinone side chains), and regulators of plant growth (the phytohormone gibberellin and phytol side chain in chlorophyll)^15^. Specialized volatile or semi-volatile terpenoids are also important biological and ecological signals that protect plants against abiotic stress and promote beneficial biotic interactions above and below ground with pollinators, pathogens, herbivorous insect, and soil microbes^15-18^. Analyses of the SCT genome inform understanding of gene family evolution contributing to terpenoid biosynthesis, shed light on early events in flowering plant diversification, and provide new insights into the demographic history of SCT with important implications for future conservation efforts.

## Results

### Assembly and annotation of SCT

SCT is diploid (2n=24; Supplementary Fig. 1a) with an estimated genome size of 800 to 846 Mb (Supplementary Figs. 1b, 2). An initial assembly with 141x and 50x Illumina paired-end and mate-pair reads, respectively (Supplementary Table 1), produced 48,650 scaffolds spanning 714.7 Mb (scaffold N50 = 594 kb and N90 = 3 kb; Table 1). A second, long-read assembly derived solely from 85x Pacbio long reads (read N50 = 11.1 kb; contig N50 = 0.9 Mb) was scaffolded with 207x “Chicago” reconstituted-chromatin and 204x Hi-C paired-end reads using the HiRise pipeline^19^ (Table 1; Supplementary Fig. 3). A final, integrated assembly of 730.7 Mb was produced in 2,153 scaffolds, comprising 91.3% of the flow cytometry genome size estimate. The final scaffold N50 was 50.4 Mb with more than 90% in 12 pseudomolecules, presumably corresponding to the 12 SCT chromosomes. Using a combination of reference plant protein homology support and transcriptome sequencing derived from a variety of tissues (Supplementary Fig. 1c and Table 2) and *ab initio* gene prediction, 27,899 protein-coding genes models were annotated using the MAKER2 pipeline^20^ (Table 1). Of these, 93.7% were found to be homologous to proteins in the TrEMBL database and 50% could be assigned gene ontology terms using eggNOG-mapper^21^. The proteome was estimated to be at least 89% complete based on BUSCO^22^ (Benchmarking Universal Single-Copy Orthologs) assessment which is comparable to other sequenced plant species (Supplementary Table 3). Orthofinder^23^ clustering of SCT gene models with those from twelve diverse seed plant genomes yielded 20,658 orthologous groups (OGs) (Supplementary Table 4). 24,148 SCT genes (85.8%) were part of OGs with orthologues from at least one other plant species. 3,744 gene models were not orthologous to others, and only 210 genes were part of the 48 SCT specific OGs. Altogether, they suggest that the phenotypic diversification in magnoliids may be fueled by *de novo* birth of species-specific genes as well as expansion of existing gene families.

**Table 1.** Statistics of stout camphor tree genome assemblies using different sequencing technologies and final gene predictions.

|  | **Illumina (v1)** | **Pacbio (v2)** | **Pacbio + HiC (v3)** |
| --- | --- | --- | --- |
| **Assembly feature** |  |  |  |
| Size (Mb) | 714.7 | 728.3 | 730.7 |
| Num. scaffolds | 48,650 | 5,210 | 2,153 |
| Average (Kb) | 15 | 140 | 339 |
| Largest scaffold (Mb) | 6.2 | 12.8 | 87.0 |
| N50 (Mb) | 0.6 | 0.9 | 50.4 |
| L50 (n) | 254 | 185 | 6 |
| N90 (Mb) | 0.003 | 0.058 | 37.1 |
| L90 (n) | 6,123 | 1,709 | 12 |
| **Genome annotation** |  |  |  |
| Num. genes (n) |  |  | 27,899 |
| Gene model (Mb) |  |  | 235.6 |
| Exons (Mb) |  |  | 36.4 |
| Intron length (Mb) |  |  | 199.2 |
| Num. genes in 12 largest scaffolds |  |  | 26,692 |
| BUSCO completeness (%) |  |  | 88.5 |

**Table 2.** Comparison of the known/predicted seven TPS subfamilies among 14 known genomes and three available transcriptomes of major seed plant lineages.

| TPS subfamilies | Genome size (Mb) | Primary Metabolism |  | Secondary Metabolism |  |  |  |  | Total no. |
| --- | --- | --- | --- | --- | --- | --- | --- | --- | --- |
|  |  | c | e | a | b | d<sup>2</sup> | f | g |  |
| Function |  | CPS, C20<sup>1</sup> | KS, C20 | C15 | IspS, C10 | C10,15,20 | C20 | C10 |  |
| Species |  |  |  |  |  |  |  |  |  |
| Gymnosperms |  |  |  |  |  |  |  |  |  |
| *Ginkgo biloba* | 10,609 | 1 | 1 | - | - | 49 | - | - | 51 |
| *Picea abies* | 12,301 | 2 | 1 | - | - | 59 | - | - | 62 |
| Angiosperms |  |  |  |  |  |  |  |  |  |
| *Amborella trichopoda* | 706 | 1 | 1 | - | 7 | - | 3 | 5 | 17 |
| Chloranthaceae |  |  |  |  |  |  |  |  |  |
| *Sarcandra glabra*<sup>3</sup> | - | - | 1 | 2 | - | - | - | - | 3 |
| Magnoliids |  |  |  |  |  |  |  |  |  |
| Lauraceae |  |  |  |  |  |  |  |  |  |
| *Cinnamomum kanehirae* | 731 | 2 | 5 | 25 | 58 | - | 7 | 4 | 101 |
| *Persea americana*<sup>3</sup> | - | - | - | 11 | 12 | - | 1 | 9 | 33 |
| Piperales |  |  |  |  |  |  |  |  |  |
| *Saruma henryi*<sup>3</sup> | - | 1 | - | 1 | 2 | - | - | 1 | 5 |
| Monocots |  |  |  |  |  |  |  |  |  |
| *Musa acuminata* | 473 | 2 | 2 | 21 | 13 | - | 3 | 3 | 44 |
| *Oryza sativa* | 375 | 3 | 10 | 19 | - | - | - | 1 | 33 |
| *Zea mays* | 2,068 | 8 | 6 | 30 | 2 | - | - | 5 | 51 |
| Eudicots |  |  |  |  |  |  |  |  |  |
| *Aquilegia coerulea* | 307 | 15 | 13 | 12 | 34 | - | - | 8 | 82 |
| Rosids |  |  |  |  |  |  |  |  |  |
| *Arabidopsis thaliana* | 120 | 1 | 1 | 23 | 5 | - | 1 | 1 | 32 |
| *Populus trichocarpa* | 473 | 2 | 2 | 16 | 14 | - | 1 | 3 | 38 |
| *Vitis vinifera*<sup>4</sup> | 434 | 2 | 1 | 29 | 10 | - | 2 | 14 | 58<sup>4</sup> |
| **Asterids** |  |  |  |  |  |  |  |  |  |
| *Daucus carota* | 422 | 3 | 2 | 1 | 15 | - | 1 | 7 | 29 |
| *Mimulus guttatus* | 313 | 13 | 13 | 19 | 17 | - | - | 1 | 63 |

<sup>1</sup> The “C” and “Arabic number” within the parenthesis designate C10: monoterpene; C15: sesquiterpenes; C20: diterpenes.
CPS: copalyl diphosphate synthase; KS: kaurene synthase; IspS: isoprene synthase.
<sup>2</sup> TPS-d subfamily is gymnosperm-specific.
<sup>3</sup> Transcriptome data of these three taxa were highly likely incomplete for covering all TPS transcripts, so that their total numbers of TPS were not reliable but for reference only.
<sup>4</sup> These two TPS-f were previously characterized from grape floral cDNA without identical genomic *VvTPS* genes (Martin et al., 2010); *VvTPS* sequences that labeled as unknown (Martin et al., 2010) in the TPS gene tree were not counted.

### Genome characterization

We identified 3,950,027 bi-allelic heterozygous sites in the SCT genome, corresponding to an average heterozygosity of 0.54% (one heterozygous SNP per 185 bp). The minor allele frequency of these sites had a major peak around 50% consistent with the fact that SCT is diploid with no evidence for recent aneuploidy (Supplementary Fig. 4). The spatial distribution of heterozygous sites was highly variable with 23.9% of the genome exhibiting less than 1 SNP loci per kb compared to 10% of the genome with at least 12.6 SNP loci per kb. Runs of homozygosity (ROH) regions appeared to be distributed randomly across SCT chromosomes reaching a maximum of 20.2 Mb in scaffold 11 (Fig. 1a). Such long ROH regions may be associated with selective sweeps, inbreeding or recent population bottlenecks. Pairwise sequentially Markovian coalescent^24^ (PSMC) analysis based on heterozygous SNP densities implicated a continuous reduction of effective population size over the last 9 Ma (Fig. 1b) with a possible bottleneck coincident with the mid-Pleistocene climatic shift at 0.9 Ma. Such patterns may reflect a complex population history of SCT associated with the geologic history of Taiwan including uplift and formation of the island in the late Miocene (9 Ma) followed by mountain building 5–6 Ma, respectively^25^.

**Figure 1.**
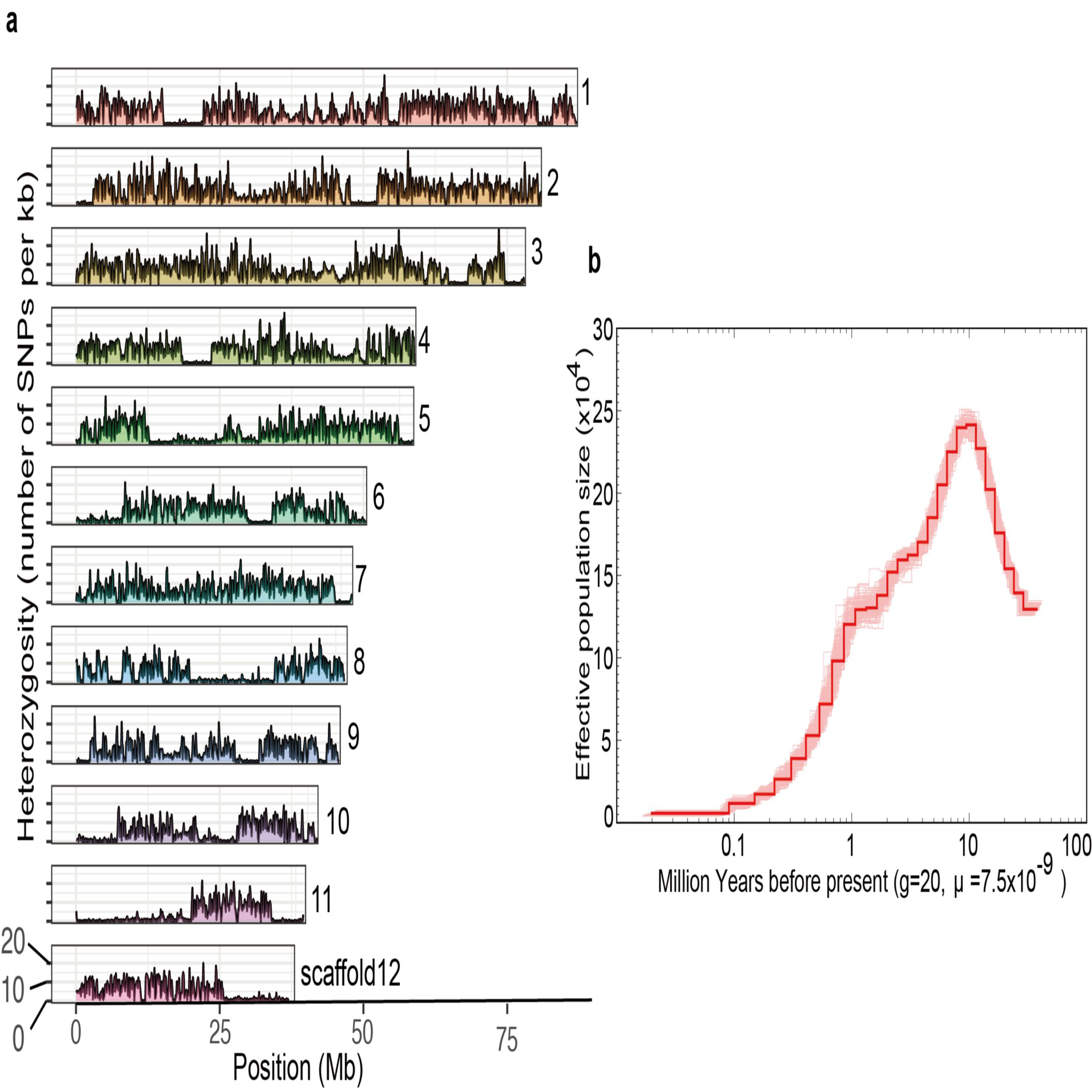
Stout camphor tree genome heterozygosity. **a,** Number of heterozygous bi-allelic SNPs per 100 kb non-overlapping windows is plotted along the largest 12 scaffolds. Indels were excluded. **b,** The history of effective population size was inferred using the PSMC method. 100 bootstraps were performed and the margins are shown in light red.

Transposable elements (TEs) and interspersed repeats made up 48% of the genome assembly (Supplementary Table 5). The majority of the TEs belonged to LTR retrotransposons (25.53%), followed by DNA transposable elements (12.67%). Among the LTR, 40.75% and 23.88% or retrotransposons belonged to Ty3/Gypsy and Ty1/Copia, respectively (Supplementary Table 5). Phylogeny of reverse transcriptase domain showed that the majority of Ty3/Gypsy copies formed a distinct clade (20,092 copies) presumably as a result of recent expansion and proliferation, while Ty1/Copia elements were grouped into two sister clades (7,229 and 2,950 copies; Supplementary Fig. 5). With the exception of two scaffolds, both Ty3/Gypsy and Ty1/Copia LTR TEs were clustered within the pericentromeric centers of the 12 largest scaffolds (Fig 2; Supplementary Fig. 6). Additionally, the LTR enriched regions (defined by 100 kb with excess of 50% comprising LTR class TEs) had on average 35% greater coverage than rest of the genome (Fig 2; Supplementary Fig. 7), suggesting that these repeats were collapsed in the assembly and may have contributed to the differences in flow cytometry and k-mer genome size estimates. The coding sequence content of SCT is similar to the other angiosperm genomes included in our analyses (Supplementary Table 3), while introns are slightly longer in SCT due to a higher density of TEs (*P* < 0.001, Wilcoxon rank sum test; Supplementary Fig. 8).

**Figure 2.**
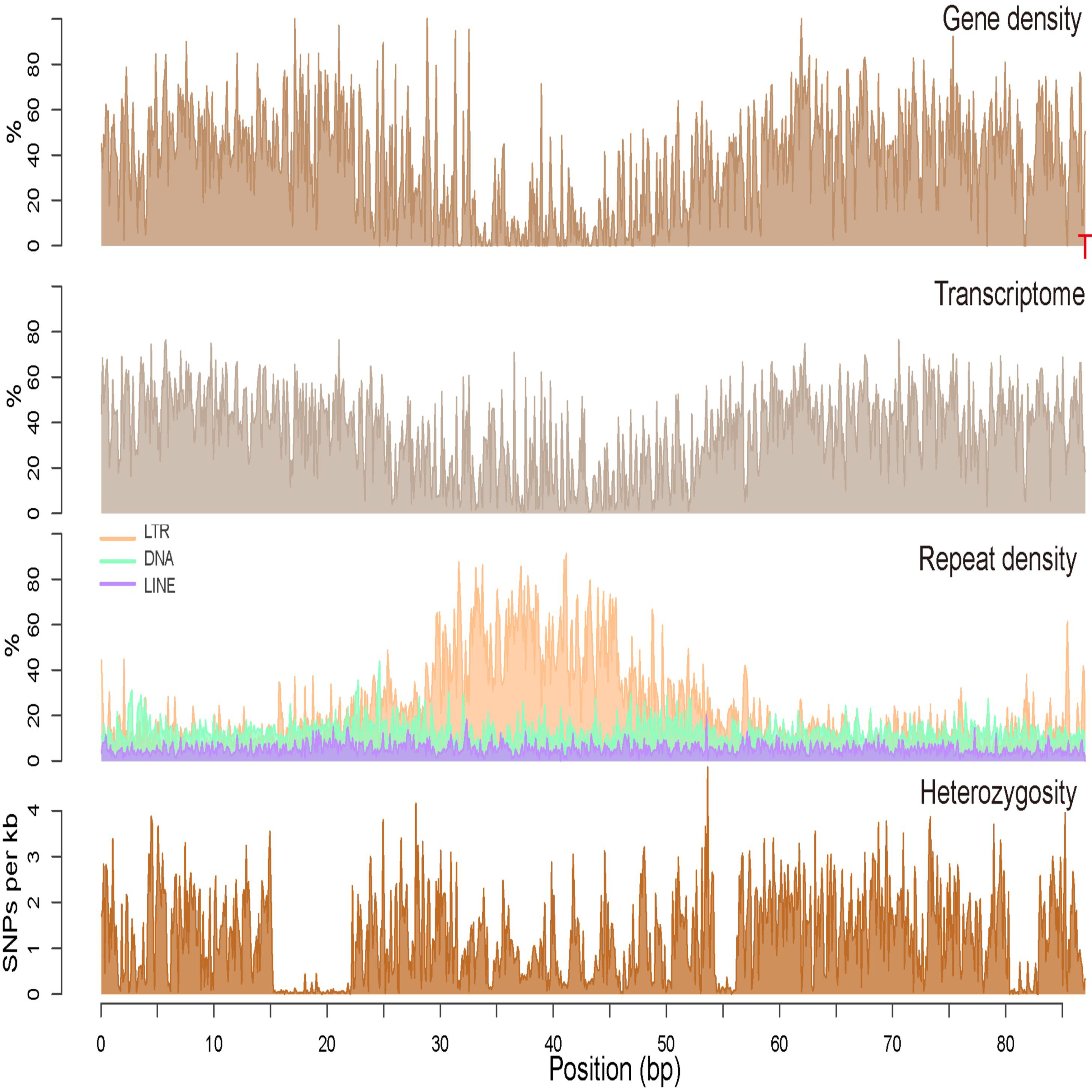
Genomic landscape of stout camphor tree chromosome 1. For every non-overlapping 100 kb window distribution is shown from top to bottom: gene density (percent of nucleotide with predicted model), transcriptome (percent of nucleotides with evidence of transcriptome mapping), three different classes repetitive sequences (percent of nucleotides with TE annotation) and heterozygosity (number of bi-allelic SNPs). The red T letter denote presence of telomeric repeat cluster at scaffold end.

As has been described for other plant genomes^26^, the chromosome-level scaffolds of SCT exhibit low protein-coding gene density and high TE density in the centers of chromosomes, and increased gene density towards the chromosome ends (Fig. 2). We identified clusters of putative subtelomere heptamer TTTAGGG extending as long as 2,547 copies, which implicate telomeric repeats in plants^27^ (Supplementary Table 6). Additionally, 687 kb of nuclear plastid DNAs (NUPT) averaging around 202.8 bp were uncovered (Supplementary Table 7). SCT NUPTs were overwhelmingly dominated by short fragments with 96% of the identified NUPTs less than 500 bp (Supplementary Table 8). The longest NUPT is ~20 kb in length and syntenic with 99.7% identity to a portion of the SCT plastome that contains seven protein-coding and five tRNA genes (Supplementary Fig. 9).

### Phylogenomic placement of *C. kanehirae* sister to eudicots

The magnoliids have been hypothesized as the sister lineage to (1) the Chloranthaceae, (2) a clade including eudicots, Chloranthaceae, Ceratophyllaceae, (3) the monocots, (4) a monocot + eudicot clade, or (5) a Chloranthaceae + Ceratophyllaceae clade, based on phylogenetic analyses of plastid genes, plastomic IR regions, four mitochondrial genes, inflorescence and floral structures, and low copy nuclear genes^7,28^. Similar to the APG III, the APG IV system^29^ placed Magnoliidae and Chloranthaceae together as sister to a robust clade comprising monocots and Ceratophyllales + eudicots. To resolve the long-standing debate over the phylogenetic placement of magnoliids relative to other major flowering plant lineages, we constructed a phylogenetic tree based on 211 strictly single copy orthologue sets shared among the 13 genomes included in our analyses. A single species tree was recovered through maximum likelihood analysis^30^ of a concatenated supermatrix of the single copy gene alignments and coalescent-based analysis using the 211 gene trees^31^ (Fig. 3; Supplementary Fig. 10). SCT, representing the magnoliid lineage was placed as sister to the eudicot clade (Fig. 3). Using MCMCtree^32^, we calculated a 95% confidence interval for the time of divergence between magnoliids and eudicots to be 139.41–191.57 million years (Ma; Supplementary Fig. 11), which overlaps with two other recent estimates (114.75–164.09 Ma^33^ and 118.9–149.9 Ma^34^).

**Figure 3.**
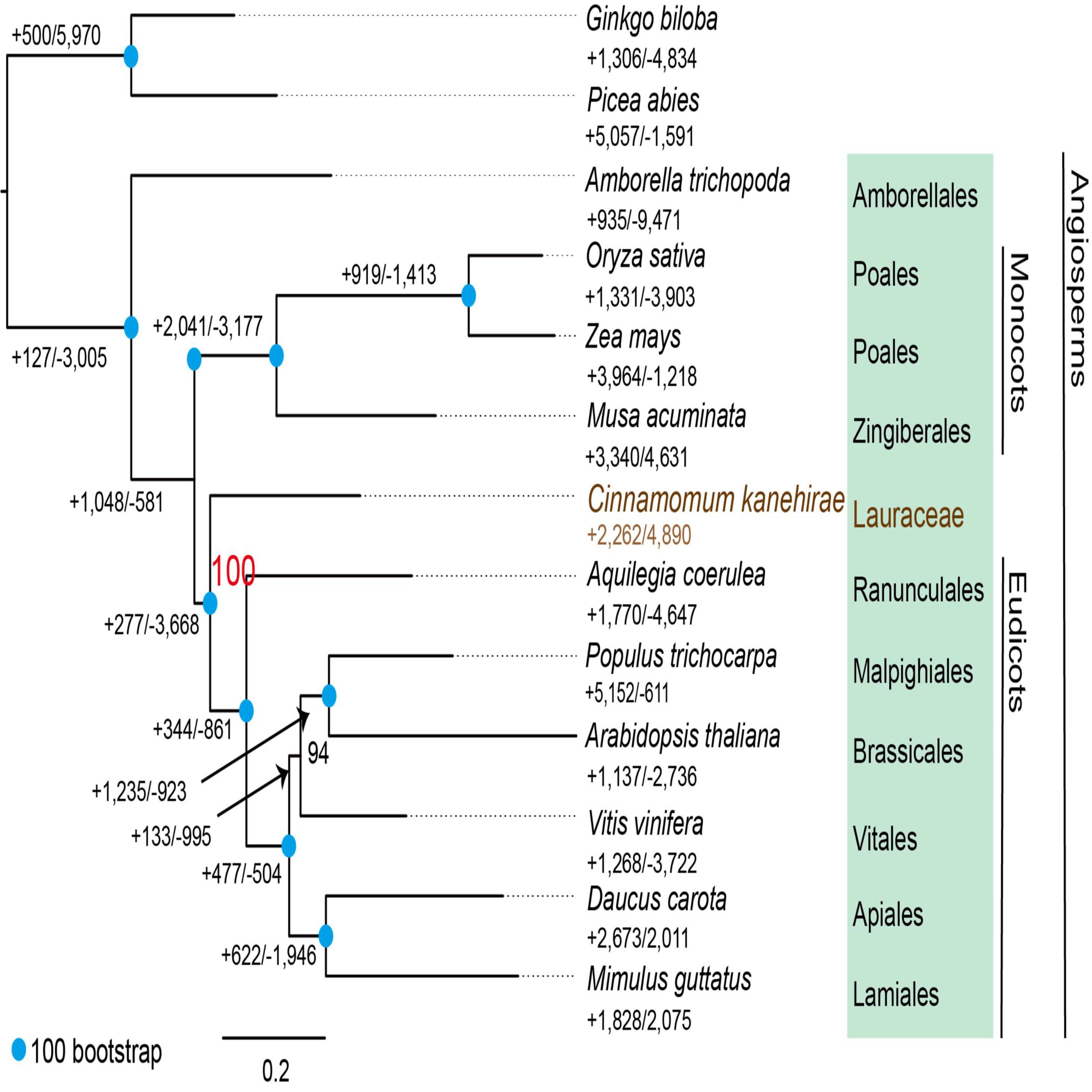
A species tree on the basis of 211 single copy orthologues from 13 plant species. Gene family expansion and contraction are denoted in numbers next to plus and minus signs, respectively. Unless stated, bootstrap support of 100 is denoted as blue circles.

### Synteny analysis / whole genome duplication (WGD)

Previous investigations of EST data inferred a genome-wide duplication within the magnoliids before the divergence of the Magnoliales and Laurales^7^, but synteny-based testing of this hypothesis has not been possible without an assembled magnoliid genome. A total of 16,498 gene pairs were identified in 992 syntenic blocks comprising 72.7% of the SCT genome assembly. Of these intragenomic syntenic blocks, 72.3% were found to be syntenic to more than one location on the genome, suggesting that more than one WGD occurred in the ancestry of SCT (Fig. 4a). Two rounds of ancient WGD were implicated by extensive synteny between pairs of chromosomal regions and significantly but less syntenic paring of each region with two additional genomic segments (Supplementary Fig. 12). Synteny blocks of SCT’s 12 largest scaffolds were assigned to five clusters that may correspond to pre-WGD ancestral chromosomes (Fig. 4a; Supplementary Fig. 12 and Note).

**Figure 4.**
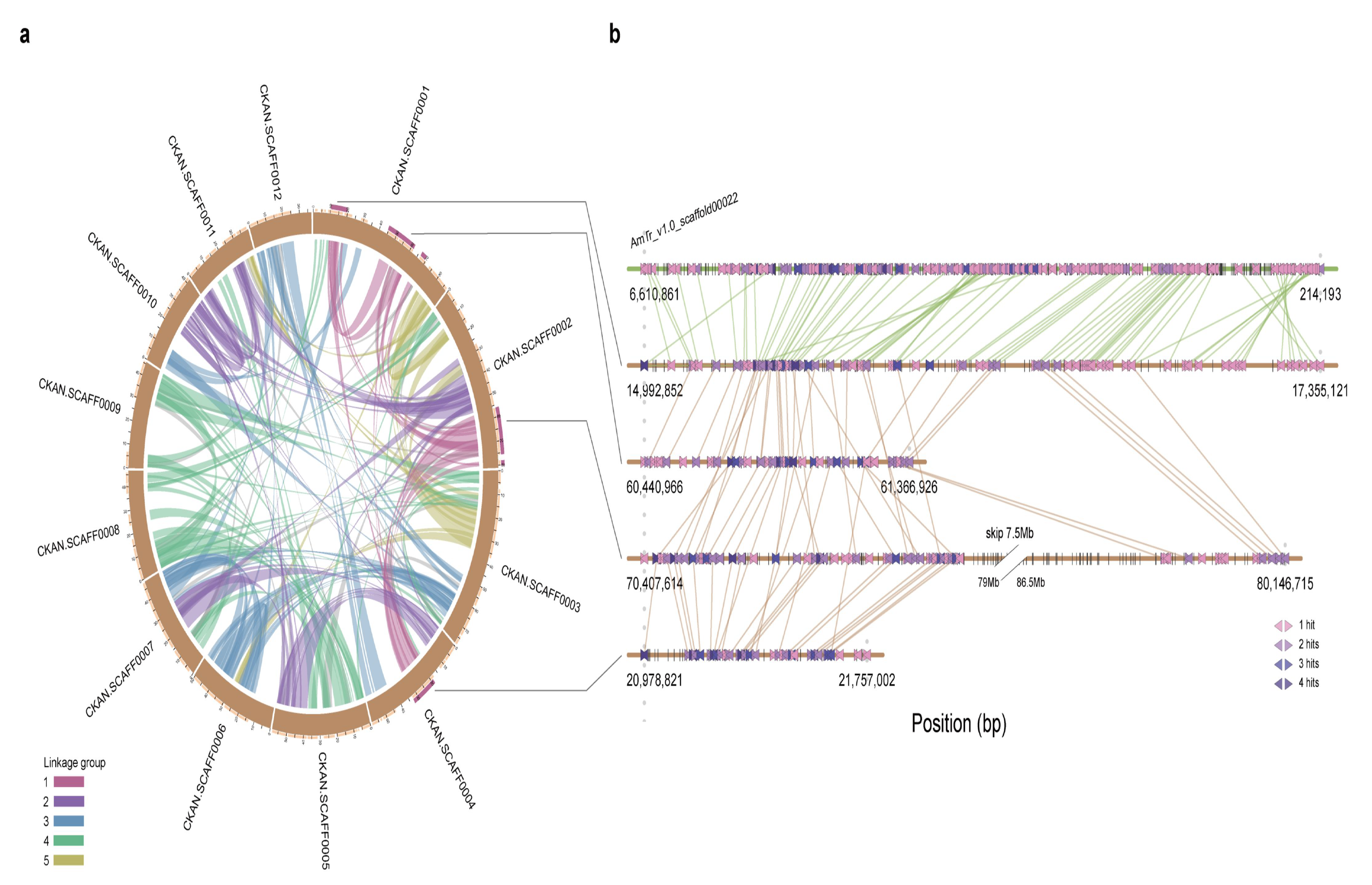
Evolutionary analysis of the stout camphor tree genome. **a,** Schematic representation of intragenomic relationship amongst the 637 synteny blocks in the stout camphor tree genome. Synteny blocks assigned unambiguously into 5 linkage clusters representing ancient karyotypes are color coded. **b,** Schematic representation of the first linkage group within the stout camphor tree genome and their corresponding relationship in *A. trichopoda*.

*Amborella trichopoda* is the sole species representing the sister lineage to all other extant angiosperms, and it has no evidence of WGD since divergence from the last common ancestor extant flowering plant lineages^35^. To confirm two rounds of WGD took place in ancestry of SCT after divergence of lineages leading to SCT and *A. trichopoda*, we assessed synteny between the two genomes. Consistent with our hypothesis, four segments of the SCT genome aligned with a single region in the *A. trichopoda* genome (Fig. 4b; Supplementary Fig. 13).

In order to more precisely infer the timing of the two rounds of WGD evident in the SCT genome, intragenomic and interspecies homolog Ks (synonymous substitutions per synonymous site) distributions were estimated. SCT intragenomic duplicates showed two peaks around 0.46 and 0.76 (Fig. 5a), congruent with the two WGD events. Based on these two peaks, we were able to infer the karyotype evolution by organizing the clustered synteny blocks further into four groups presumably originating from one of the five pre-WGD chromosomes (Supplementary Fig. 14). Comparison between *Aquilegia coerulea* (Ranunculales, a sister lineage to all other extant eudicots^35^) and SCT orthologs revealed a prominent peak around Ks = 1.41 (Fig. 5a), while the *Aquilegia* intra-genomic duplicate was around Ks = 1, implicating independent WGDs following the divergence of lineages leading to SCT and *Aquilegia.* The availability of the transcriptome of 17 Laurales + Magnoliales from 1,000 plants initiative^36^ allowed us to test the hypothesized timing of the WGDs evident in the SCT genome^8^. Ks distribution of all species from Lauraceae have shown apparent two peaks, but only one peak was observed in other Laurales and Magnoliales samples, suggesting a WGD predating divergence of these two orders followed by a second recent WGD in the early ancestry of the Lauraceae (Fig. 5b). The Ks peak seen in *Aquilegia* data is likely attributable to WGD within the Ranunculales well after the divergence of eudicots and magnoliids (Supplementary Fig. 15).

**Figure 5.**
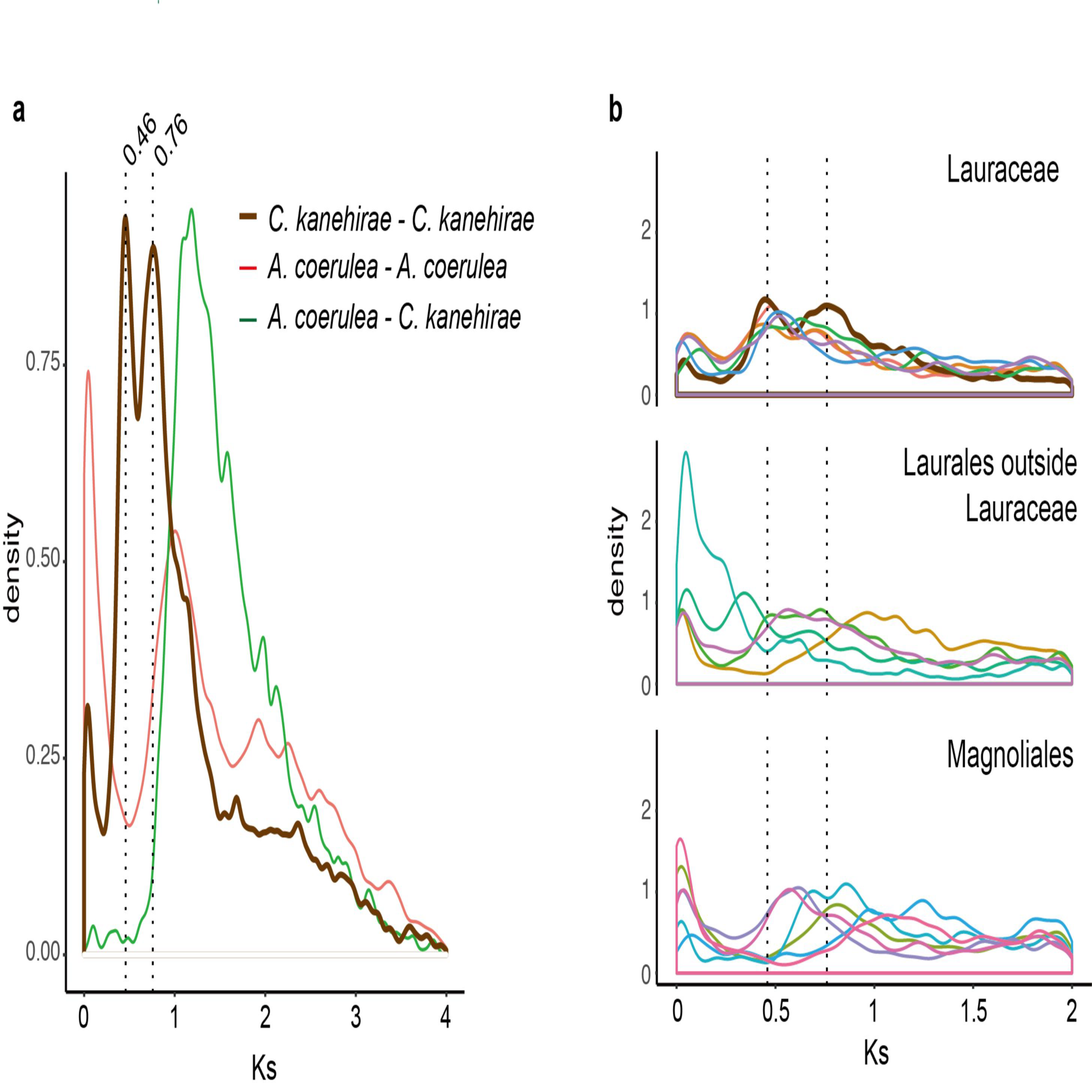
Density plots of synonymous substitutions (Ks) of stout camphor tree genome and other plant species. **a,** Pairwise orthologue duplicates identified in synteny blocks within SCT, within *A. coerulea* and between SCT and *A. coerulea*. **b,** Ks of intragenomic pairwise duplicates of the Lauraceae and the Magnoliales in the 1KP project^104^. Dashed lines denote the two Ks peaks observed in SCT.

### Specialization of the magnoliids proteome

We sought to identify genes and protein domains specific to SCT by annotating protein family (Pfam) domains^37^ and assessing their distribution across the 13 seed plant genomes included in our phylogenomic analyses. Consistent with the observation that there were very few SCT-specific OGs, principal component analysis of Pfam domain content clustered SCT with the monocots and eudicots, with the first two principal components separating gymnosperms and *A. trichopoda* from this group (Supplementary Fig. 16a). There were considerable overlaps between SCT, eudicot and monocot species, suggesting significant functional diversification since these three lineages split. SCT also showed a significant enrichment and reduction of 111 and 34 protein domains compared to other plant species, respectively (Supplementary Fig. 16b and Table 9). Gain of protein domains included the terpene synthase C terminal domain involved in defense responses and the leucine-rich repeats (628 vs 334.4) in plant transpiration efficiency^38^. Interestingly, we found that SCT possesses 21 copies of EIN3/EIN3-like (EIL) transcription factor, more than the previously reported maximum of 17 copies in the banana genome (*Musa acuminata*)^39^. EILs initiate an ethylene signaling response by activating ethylene response factors (ERF), which we also found to be highly expanded in SCT (150 copies versus an average of 68.3 copies from nine species reported in ref^39^; Supplementary Fig. 17). Ethylene signaling in plants was reported to be associated with fruit ripening^39^ and secondary growth in wood formation^40^ and may be involved in either processes in SCT.

CAFE^41^ was used to assess OG expansions and contractions across (Fig. 3) the seed plant phylogeny. Gene family size evolution was dynamic across the phylogeny, and the branch leading to SCT did not exhibit significantly different numbers of expansions and contractions. Enrichment of gene ontology terms revealed either various different gene families sharing common functions or single gene families undergoing large expansions (Supplementary Table 10 and 11). For example, the expanded members of plant resistance (R) genes add up to “plant-type hypersensitive response” (Supplementary Table 10). In contrast, the enriched gene ontology terms from the contracted gene families of SCT branch (Supplementary Table 11) contains members of ABC transporters, indole-3-acetic acid-amido synthetase, xyloglucan endotransglucosylase/hydrolase and auxin-responsive protein, all of which are part of the “response to auxin”.

### Resistance (R) genes

The SCT genome annotation included 387 resistance gene models, 82% of which belong to nucleotide-binding site leucine-rich repeat (NBS-LRR) or coiled-coil NBS-LRR (CC-NBS-LRR) types. This result is consistent with a previous report that LRR is one of the most abundant protein domains in plants and it is highly likely that SCT is able to recognize and fight off pathogen products of avirulence (Avr) genes^42^. Among the sampled 13 genomes, SCT harbors the highest number of R genes among non-cultivated plants (Supplementary Fig. 18). The phylogenetic tree constructed from 2,465 NBS domains also suggested that clades within the gene family have diversified independently within the eudicots, monocots and magnoliids. Interestingly, the most diverse SCT NBS gene clades were sister to depauperate eudicot NBS gene clades (Supplementary Fig. 19).

### Terpene synthase gene family

One of the most striking features of the SCT genome is the large number of terpene synthase (TPS) genes (*CkTPSs*). A total of 101 *CkTPSs* were predicted and annotated, the largest number for any other genome to date. By including transcriptome dataset of two more species from magnoliids (*Persea americana* and *Saruma henryi*), phylogenetic analyses of TPS from 15 species were performed to place *CkTPSs* among six of seven TPS subfamilies that have been described for seed plants^43-45^ (Fig. 6, Table 2 and Supplementary Fig. 20–25). *CkTPS* genes placed in the TPS-c (2) and TPS-e (5) subfamilies likely encode diterpene synthases such as copalyl diphosphate synthase (CPS) and *ent*-kaurene synthase (KS)^46^. These are key enzymes catalyzing the formation of the 20-carbon isoprenoids (collectively termed diterpenoids; C20), which was thought to be eudicot-specific^45^ and serve primary functions like regulating plant primary metabolism. The remaining 94 predicted *CkTPSs* likely code for the 10-carbon monoterpene (C10) synthases, 15-carbon sesquiterpene (C15) synthases, and additional 20-carbon diterpene (C20) synthases (Table 2). With 25 and 58 homologs, respectively, TPS-a and TPS-b subfamilies are most diverse in SCT, presumably contributing to the mass and mixed production of volatile C15s and C10s^47^. *CkTPSs* are not uniformly distributed throughout the chromosomes (Supplementary Table 12) and clustering of members from individual subfamilies were observed as tandem duplicates (Supplementary Fig. 26). For instance, scaffold 7 contains 29 *Ck*TPS genes belonging to several subfamilies including all of the eight *CkTPS-a*, 12 *CkTPS-b*, five *CkTPS-e* and three *CkTPS-f* (Supplementary Fig. 26). In contrast, only two members of *CkTPS-c* reside in scaffold 1. Twenty-four *CkTPSs* locate in other smaller scaffolds, 22 of which code for subfamily TPS-b (Supplementary Fig. 21).

**Figure 6.**
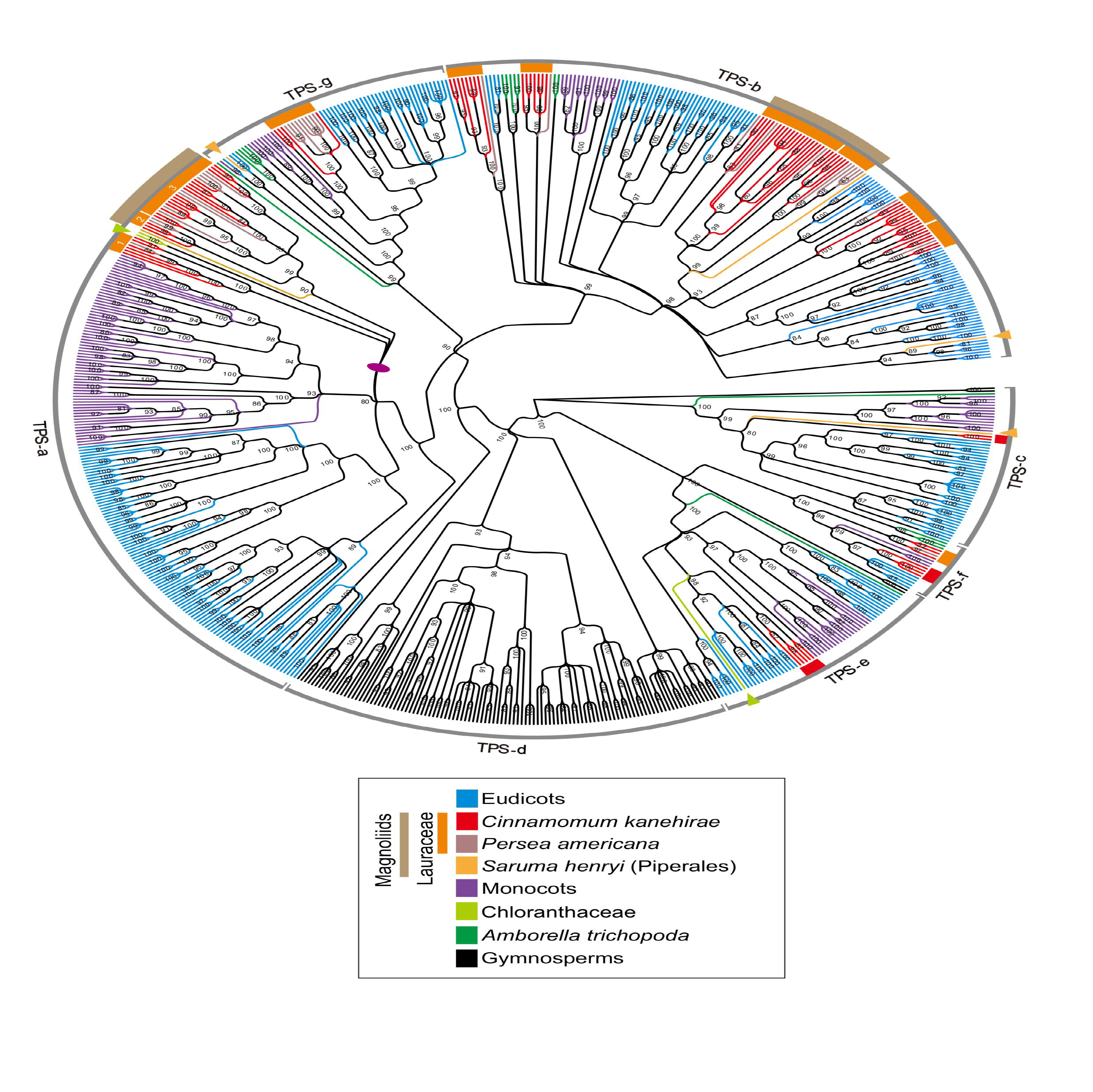
Phylogenetic tree of putative or characterized TPS genes from the 13 sequenced land plant genomes and two magnoliids with available transcriptomic data.

It is noteworthy that the TPS gene tree resolved Lauraceae-specific TPS gene clades within the TPS-a, -b, -f, and -g subfamilies (Supplementary Fig. 20–23). This pattern of TPS gene duplication in a common ancestor of *Persea* and *Cinnamomum* and subsequent retention may indicate subfunctionalization or neofunctionalization of duplicated *TPSs* within the Lauraceae. A magnoliids-specific subclade in the TPS-a subfamily was also identified in analyses including more magnoliid TPS genes with characterized functions (Supplementary Fig. 20). Indeed, we detected positive selection in the Lauraceae-specific TPS-f -I and -II subclades implying functional divergence (Supplementary Table 13). Together, these data suggest increasing diversification of magnoliid TPS genes both before and after the origin of the Lauraceae. The distribution of TPS genes in the SCT genome suggests that both segmental (including WGD) and tandem duplication events contributed to diversification of TPS enzymes in the SCT lineage and the terpenoids they produce.

## Discussion

It is now challenging to find a wild SCT population making the conservation and basic study of this tree a priority. SCTs have been intensively logged since the 19^th^ century initially for hardwood properties and association with fungus *Antrodia cinnamomea*. The apparent runs of homozygosity have been observed due to anthropogenic selective pressures or inbreeding in several livestock^47^, though inbreeding as a result of recent population bottleneck may be a more likely explanation for SCT. Interestingly, continuous decline in effective population size was inferred since 9 Ma. These observations may reflect a complex population history of SCT and Taiwan itself after origination and mountain building of the island that occurred around late Miocene (9 Ma) and 5−6 Ma, respectively^25^. The availability of the SCT genome will help the development of precise genetic monitoring and tree management for the survival of SCT’s natural populations.

The placement of SCT as sister to the eudicots has important implications for comparative genomic analyses of evolutionary innovations within the eudicots, which comprise ca. 75% of extant flowering plants^48^. For example, the SCT genome will serve as an important reference outgroup for reconstructing the timing and nature of polyploidy event that gave rise to the hexaploid ancestor of all core eudicots (Pentapetalae)^49,50^. Within the magnoliids we identified the timing of two independent rounds of WGD events that contributed to gene family expansions and innovations in pathogen, herbivore and mutualistic interactions.

Gene tree topologies for each of the six angiosperm TPS subfamilies revealed diversification of TPS genes and gene function in the ancestry of SCT. The C20s producing TPS-f genes were suggested to be eudicot-specific because both rice and sorghum lack genes in this subfamily^45^. Our data clearly indicate that this subfamily was present in the last common ancestor of all but was lost from the grass family (Table 2). Massive diversification of the TPS-a and TPS-b subfamilies within the Lauraceae is consistent with a previous report that the main constituents of 58 essential oils produced in *Cinnamomum* leaves are C10s and C15s^47^. These findings are in congruent with the fact that fruiting bodies of the SCT-specific parasitic fungus, *Antrodia cinnamomea*, can produce 78 kinds of terpenoids, including 31 structure-different triterpenoids (C30s)^51^, many of which are synthesized via the mevalonate pathway as are C10s and C15s followed by cyclizing squalenes (C_30_H_50_) into the skeletons of C30s^52^. It is reasonable to suggest that this fungus obtained intermediate compounds through decomposing trunk matters from SCT.

The 101 *CkTPSs* identified in the SCT genome are unevenly distributed across the 12 chromosomal scaffolds, and tandem arrays include gene clusters from the same subfamily (Supplementary Fig. 26). In the *Drosophila melanogaster* genome, “tandem duplicate overactivity” has been observed with tandemly duplicated *Adh* genes showing 2.6-fold greater expression than single copy *Adh* genes^53^.

In summary, the availability of SCT genome establishes a valuable genomic foundation that will help unravel the genetic diversity and evolution of other magnoliids, and a better understanding of flowering plant genome evolution and diversification. At the same time, the reference-quality SCT genome sequence will enable efforts to conserve genome-wide genetic diversity in this culturally and economically important tree species.

## Methods

### Plant Materials

All plant materials used in this study were collected from a 12-year-old SCT growing in Ershui Township, Changhua County, Taiwan (23°49′25.9”N,120°36′41.2”E) during April to July of 2014–2016. The tree was grown up from a seedling obtained from Forestry Management Section, Department of Agriculture, Taoyuan City. The specimen (voucher number: Chaw 1501) was deposited in the Herbarium of Biodiversity Research Center, Academia Sinica, Taipei, Taiwan (HAST).

### Genomic DNA extraction and sequencing

We used a modified high-salt method^54^ to eliminate the high content of polysaccharides in SCT leaves, followed by total DNA extraction with a modified CTAB method^55^. Three approaches were employed in DNA sequencing. First, paired-end and mate-pair libraries were constructed using the Illumina TruSeq DNA HT Sample Prep Kit and Illumina Nextera Mate Pair Sample Prep Kit following the kit’s instructions, respectively. All obtained libraries were sequenced on an Illumina NextSeq 500 platform to generate ca. 278.8 Gb of raw data. Second, SMRT libraries were constructed using the PacBio 20-Kb protocol (https://www.pacb.com/). After loading on SMRT cells (SMRT™ Cell 8 Pac), these libraries were sequenced on a PacBio RS-II instrument using P6 polymerase and C4 sequencing reagent (Pacific Biosciences, Menlo Park, California). Third, a Chicago library was prepared by Dovetail Genomics (Santa Cruz, California and sequenced on an Illumina HiSeq 2500 to generate 150 bp read pairs. Supplementary Table 1 summarizes the coverage and information for the sequencing data.

### RNA extraction and sequencing

Opening flowers, flower buds (two stages), immature leaves, young leaves, mature leaves, young stems, and fruits were collected from the same individual (Supplementary Fig. 1c) and their total RNAs were extracted^56^. The extracted RNA was purified using poly-T oligo-attached magnetic beads. All transcriptome libraries were constructed using Illumina TruSeq library Stranded mRNA Prep Kit and sequenced on an Illumina HiSeq 2000 platform. A summary of transcriptome data is shown in Supplementary Table 2.

### Chromosome number assessment

Root tips from cutting seedlings were used to examine the chromosome number based on Suen *et al*.’s method^57^. The stained samples were observed under a Nikon Eclipse 90i microscope (Supplementary Fig. 1a).

### Genome size estimation

Fresh leaves of SCT were cut into tiny pieces and mixed well with 1 mL isolation buffer (200 mM Tris, 4 mM MgCl_2_-6H_2_O, and 0.5% Triton X-100)^58^. The mixture was filtered through a 42 μm nylon mesh, followed by incubation of the filtered suspensions with a DNA fluorochrome (50 μg/ml propidium iodide and 50 μg/ml RNase). The genome size was estimated using a MoFlo XDP flow cytometry (Beckman Coulter Life Science, Indianapolis, Indiana) with chicken erythrocyte and rice nuclei (BioSure, Grass Valley, California) as the internal standards (Supplementary Fig. 1b). Estimate of genome size from Illumina paired end sequences was inferred using Genomescope^59^ (based on k-mer 31).

### *De novo* assembly of SCT

Illumina paired end and mate pair reads were trimmed with Trimmomatic^60^ (ver. 0.32; options LEADING:30 TRAILING:30 SLIDINGWINDOW:4:30 MINLEN:50) and subsequently assembled using Platanus^61^. Pacbio reads were assembled using the FALCON^62^ assembler and the consensus sequences were improved using Quiver^63^. The Pacbio assembly was scaffolded using HiRISE scaffolder and consensus sequences were further improved using Pilon with one iteration^64^. The genome completeness was assessed using plant dataset of BUSCO^22^ (ver. 3.0.2). To identify putative telomeric repeats, the assembly was searched for high copy number repeats less than 10 base pairs using tandem repeat finder^65^ (ver. 4.09; options: 2 7 7 80 10 50 500). The heptamer TTTAGGG was identified (Supplementary Table 6).

### Gene predictions and functional annotation

Transcriptome paired end reads were aligned to the genome using STAR^66^. Transcripts were identified using two approaches: i) assembled *de novo* using Trinity^67^, ii) reconstructed using Stringtie^68^ or CLASS2^69^. Transcripts generated from Trinity were remapped to the reference using GMAP^70^. The three sets of transcripts were merged and filtered using MIKADO (https://github.com/lucventurini/mikado). Proteomes from representative reference species (Uniprot plants; Proteomes of *Amborella trichopoda* and *Arabidopsis thaliana*) were downloaded from Phytozome (ver. 12.1; https://phytozome.jgi.doe.gov/)). The gene predictor Augustus^71^ (ver. 3.2.1) and SNAP^72^ were trained either on the gene models data using BRAKER1^73^ or MAKER2^20^. The assembled transcripts, reference proteomes, BRAKER1 and the BUSCO predictions were combined as evidence hints for input of the MAKER2^20^ annotation pipeline. MAKER2^20^ invoked the two trained gene predictors to generate a final set of gene annotation. Amino acid sequences of the proteome were functionally annotated using Blast2GO^74^ and eggnog-mapper^21^. Nuclear plastid DNAs (NUPT) of SCT was searched against its plastid genome (plastome; KR014245^75^) using blastn (parameters were followed from ref^76^).

### Analysis of genome heterozygosity

Paired end reads of SCT was aligned to reference using bwa mem^77^ (ver. 0.7.17-r1188). PCR duplicates were removed using samtools^78^ (ver. 1.8). Heterozygous bi-allelic SNPs were called using samtools^78^ and consensus sequences were generated using bcftools^79^ (ver. 1.7). Depth of coverage and minor allele frequency plots were conducted using R ver. 3.4.2. Consensus sequence was fed to the PSMC program^24^ to infer past effective population size. All of the parameters used for the PSMC program were at default with the exception of -u 7.5e-09 taken from *A. thaliana*^80^ and -g 20 taken from *Neolitsea sericea* (Lauraceae)^81^.

### Identification of repetitive elements

Repetitive elements were firstly identified by modeling the repeats using RepeatModeler^82^ and then searched and quantified repeats using RepeatMasker^83^. Repeat types modeled as “Unknown” by RepeatModeler were further annotated using TEclass^84^. Tandem Repeats were identified using Tandem Repeats Finder^65^. The proportions of different types of repeats were quantified by dissecting the 12 largest scaffolds into 100,000 bp chunks and calculating the total lengths and percentages of the repetitive elements within the chunks. LTR-RT domains were extracted following Guan *et al.*’s method^85^. Briefly, a two-step procedure was applied on the genomes. The first was to find candidate LTR-RTs similar to known reverse transcriptase domains and second was to identify other LTR-RTs using the candidates identified in the first step. The identified LTR-RT domains were integrated with those downloaded from the Ty1/Copia and Ty3/Gypsy trees of Guan *et al.*^85^. Trees were built by aligning the sequences using MAFFT^87^ (ver. 7.310; –genafpair –ep 0) and applied FastTree^88^ with JTT model on the aligned sequences, and were colored using APE package^89^.

### Gene family / Orthogroup inference and analysis of protein domains

The amino acid and nucleotide sequences of 12 representative plant species were downloaded from various sources: *Aquilegia coerulea, Arabidopsis thaliana, Daucus carota, Mimulus guttatus, Musa acuminata, Oryza sativa japonica, Populus trichocarpa, Vitis vinifera* and *Zea mays* from Phytozome (ver. 12.1; https://phytozome.jgi.doe.gov/), *Picea abies* from the Plant Genome Integrative Explorer Resource^90^ (http://plantgenie.org/), *Ginkgo biloba* from GigaDB^91^, and *Amborella trichopoda* from Ensembl plants^92^ (Release 39; https://plants.ensembl.org/index.html). Gene families or orthologous groups of these species and SCT were determined by OrthoFinder^23^ (ver. 2.2.0). Protein family domains (Pfam) of each species were calculated from Pfam website (ver. 31.0; https://pfam.xfam.org/). Pfam numbers of every species were transformed into z-scores. Significant expansion or reduction of Pfams in SCT were based on its z-score greater than 1.96 or less than −1.96, respectively. The significant Pfams were sorted by Pfam numbers (Supplementary Fig. 16). Gene family expansion and loss were inferred using CAFE^41^ (ver. 4.1 with input tree as the species tree inferred from the single copy orthologues).

### Phylogenetic analysis

MAFFT^87^ (ver. 7.271; option –maxiterate 1000) was used to align 13 sets of amino acid sequences of 211 single-copy OGs. Each OG alignment was used to compute a maximum likelihood phylogeny using RAxML^30^ (ver. 8.2.11; options: -m PROTGAMMAILGF -f a) with 500 bootstrap replicates. The best phylogeny and bootstrap replicates for each gene were used to infer a consensus species tree using ASTRAL-III^31^. A maximum likelihood phylogeny was constructed with the concatenated amino acid alignments of the single copy OGs (ver. 8.2.11; options: -m PROTGAMMAILGF -f a) also with 500 bootstrap replicates.

### Estimation of divergence time

Divergence time of each tree node was inferred using MCMCtree of PAML^32^ package (ver. 4.9g; options: correlated molecular clock, JC69 model and rest being default). The final species tree and the concatenated translated nucleotide alignments of 211 single-copy-orthologs were used as input of MCMCtree. The phylogeny was calibrated using various fossil records or molecular divergence estimate by placing soft bounds at split node of: i) *A. thaliana*-*V. vinifera* (115–105 Ma)^93^, ii) *M. acuminata-Z. mays* (115–90 Ma)^93^, iii) Ranunculales (128.63–119.6 Ma)^34^, iv) Angiospermae (247.2–125 Ma)^34^, v) Acrogymnospermae (365.629–308.14 Ma)^34^, and v) a hard bound of 420 Ma of outgroup *P. patens*^94^.

### Analysis of genome synteny and whole genome duplication

Dot plots between SCT and *A. trichopoda* assemblies were produced using SynMap from Comparative Genomics Platform (Coge^95^) to visualize the paleoploidy level of SCT. Synteny blocks within SCT and between *A. trichopoda* and *A. coerulea* were identified using DAGchainer^96^ (same parameters as Coge^95^: -E 0.05 -D 20 -g 10 -A 5). Ks between syntenic group pairs were calculated using the DECIPHER^97^ package in R. Depth of the inferred syntenic blocks were calculated using Bedtools^98^. Both the Ks distribution and syntenic block depth were used to determine the paleopolyploidy level^99^ of SCT. Using the quadruplicate or triplicate orthologues in the syntenic blocks as backbones, as well as *A. trichopoda* regions showing up to four syntenic regions, we identified the start and end coordinates of linkage clusters (Supplementary Note).

### Resistance (R) genes

R genes were identified based on the ref^100^. Briefly, the predicted genes of the 13 sampled species were searched for the Pfam NBS (NB-ARC) protein family (PF00931) using HMMER ver. 3.1b2^101^ with an e-value cutoff of 1e-5. Extracted sequences were then checked for protein domains using InterproScan^102^ (ver. 5.19-58.0) to remove false positive NB-ARC domain hits. The NBS domains of the genes that passed both HMMER and InterproScan were extracted according to the InterproScan annotation and aligned using MAFFT^87^ (ver. 7.310; –genafpair –ep 0); the alignment was then input into FastTree^88^ with the JTT model and visualized using EvolView^103^.

### Terpene synthase genes

In addition to the 13 species’ proteome dataset used in this study, transcriptome data from one Chloranthaceae species, *Sarcandra glabra* and two magnollids representatives, *Persea americana* (avocado) and *Saruma henryi* (saruma), were downloaded from oneKP transcriptome database^104^. Previously annotated TPS genes of four species: *Arabidopsis thaliana^105^*, *Oryza sativa*^45^, *Populus trichocarpa*^106^, and *Vitis vinifera^107^* were retrieved. For species without *a priori* TPS annotations, two Pfam domains: PF03936 and PF01397, were used to identify against the proteomes using HMMER^108^ (ver. 3.0; cut-off at e-values < 10^-5^). Sequence lengths shorter than 200 amino acids were excluded from further analysis. 702 putative or annotated protein sequences of *TPS* were aligned using MAFFT^87^ (ver. 7.310 with default parameters) and manually adjusted using MEGA^109^ (ver. 7.0). The TPS gene tree was constructed using FastTree^110^ (ver. 2.1.0) with 1,000 bootstrap replicates. Subfamily TPS-c was designated as the outgroup. Branching nodes with bootstrap values < 80% were treated as collapsed.

## Authors contribution

Conceived the study: S.M.C

Genome assembly and annotation: I.J.T and H.M.K

Repeat Analysis: L.Y.C and Y.W.W

Plastid DNA analysis: E.S.

Conducted the experiments: C.S.W, L.N.W, H.T.Y., C.Y.H and S.M.C.

Comparative genomics analysis: I.J.T, Y.L, H.M.K, C.Y.I.L and J.L.M

Analysis of R genes: Y.W.W, M.H.H, K.P.W, S.M.C

Analysis of terpene gene family: H.Y.W, S.M.C, C.Y.H and Y.W.W

Wrote the manuscript: I.J.T, J.LM and S.M.C

## Data availability

All of raw sequence reads used in this study have been deposited in NCBI under the BioProject accession number PRJNA477266. The assembly and annotation of SCT is available under the accession number SAMN09509728.

## Acknowledgement

Chi-yuan Tsai for plant materials; Chih-Ming Hung for PSMC analysis. S.M.C was funded by Investigators’ Award and Central Academic Committee, Academia Sinica. I.J.T was funded by Career Development Award, Academia Sinica. H.M.K, C.S.W and C.Y.H were funded by postdoctoral fellowship, Academia Sinica.

